# Slow conformational exchange and overall rocking motion in ubiquitin protein crystals

**DOI:** 10.1101/126813

**Authors:** Vilius Kurauskas, Sergei A. Izmailov, Olga N. Rogacheva, Audrey Hessel, Isabel Ayala, Joyce Woodhouse, Anastasya Shilova, Yi Xue, Tairan Yuwen, Nicolas Coquelle, Jacques-Philippe Colletier, Nikolai R. Skrynnikov, Paul Schanda

## Abstract

Proteins perform their functions in solution but their structures are most frequently studied inside crystals. Here we probe how the crystal packing alters microsecond dynamics, using solid-state NMR measurements and multi-microsecond MD simulations of different crystal forms of ubiquitin. In particular, NEar-Rotary-resonance Relaxation Dispersion (NERRD) experiments probe angular backbone motion, while Bloch-McConnell Relaxation Dispersion data report on fluctuations of the local electronic environment. These experiments and simulations reveal that the packing of the protein can significantly alter the thermodynamics and kinetics of local conformational exchange. Moreover, we report small-amplitude reorientational motion of protein molecules in the crystal lattice with a ∼3-5° amplitude on a tens-of-microseconds time scale in one of the crystals, but not in others. An intriguing possibility arises that overall motion is to some extent coupled to local dynamics. Our study highlights the importance of considering the packing when analyzing dynamics of crystalline proteins.

## Introduction

Proteins perform their functions in aqueous solution at ambient temperatures by sampling a multitude of conformational states involved, for example, in enzymatic turnover or binding to different partners. Understanding protein function requires characterizing the three-dimensional structures, ideally of all these thermally accessible conformations, and deciphering how the different conformations interconvert. The structural basis for understanding protein function have been laid primarily through X-ray diffraction (XRD) of proteins embedded in crystals. Recent methodological advances in XRD aim at characterizing also the structural heterogeneity by fitting multiple conformers – rather than a single structure – into the electron density obtained from Bragg diffraction peaks, or by additionally making use of diffuse scattering intensity. ^1-6^ The direct structural insight into protein dynamics that one may obtain from such ensemble crystallographic approaches may shed light on molecular mechanisms of functional processes; in particular, when combined with techniques that provide access to kinetics and thermodynamics, such as solution-state NMR spectroscopy^7^, one may obtain a fairly complete spatio-temporal picture, which may ultimately lead to a more detailed understanding of protein function.

The densely packed environment of crystals differs substantially from solution, raising the question how packing may alter the properties of proteins. Structures obtained from crystalline proteins are generally in excellent agreement with state-of-the-art solution-NMR structures, and are generally as good in predicting solution-NMR structural parameters as the NMR structures themselves.^8,9^ However, whether dynamics, i.e. the ensemble of co-existing conformers and their pairwise interconversion rates, are also conserved in crystals is less well studied. Evidence that the dynamics required for function are retained in crystals, at least in some cases, comes from the observation of functional activity, such as ligand binding and enzymatic catalysis, in crystals.^10-14^ Direct experimental investigations of the dynamics of proteins in crystals at the atomic level have turned out to be difficult. Early studies focused on comparing XRD temperature factors (B-factors) with solution-state NMR order parameters (*S*^2^).^15-17^ Such comparisons only had limited success and remained ambiguous, because of the very different nature and information content of these observables (see ref. 18 and references therein). Recent developments in magic-angle spinning solid-state NMR (MAS ssNMR) of proteins have allowed a more direct comparison of NMR parameters in solution and crystals.^19-21^ Furthermore, comparative molecular dynamics (MD) simulations of proteins embedded in an explicit crystal lattice and in solution also produced useful insights, at least over the time scales accessible to MD (typically sub-microsecond).^22-25^ The picture arising from these NMR and MD studies suggests that the crystal lattice has indeed only little impact on dynamics on sub-microsecond time scales: changes are primarily found in loops engaged in crystal contacts, whereas secondary structure elements and loops remote from neighboring molecules showed similar order parameters. Many biological processes, however, occur on longer time scales, often micro-to-milliseconds (μs-ms), and they may involve rare excursions to low-populated “minor states”. In many cases, such rare conformations seem to be crucial for functional activity,^26^ and they may involve larger-scale rearrangements comprising a bigger portion of the structure. Whether such slow dynamic processes are sustained in crystals, and how the packing alters these motions is poorly understood. MD simulation methods only recently attained the ability to model those time scales, and experimental NMR methods likewise became available only recently.

Here we use a combination of ssNMR experimental methods with microsecond-long MD simulations of explicit crystal lattices, in order to investigate how the crystalline environment impacts slow dynamics of the 8.6 kDa regulatory protein ubiquitin. The dynamics of this protein in solution have been extensively characterized experimentally and computationally. In particular, microsecond motion has been shown for the loop connecting ubiquitin’s β4 strand and the 3_10_ helix, comprising residues E51 to R54, which exists in two distinct states that differ by a flip of the D52/G53 peptide plane, along with an alteration of hydrogen bonds within this β-turn and to the adjacent helix.^27-32^ These two states correspond to so-called type-I and type-II β-turns, respectively, herein referred to as βI and βII states. The dynamic exchange process between these two conformations has been proposed to result in small (within 1 Å) but discernible reshaping of the entire molecule,^30^ and perturbation of the β-turn equilibrium has been shown to allosterically alter the binding of proteins to a surface located on the opposite side of the ubiquitin molecule.^30^ This dynamic process is thus thought to be of considerable functional relevance for ubiquitin. Different crystal structures are available that are in either of these two states, and millisecond-long MD simulations in solution have detected the corresponding microsecond exchange process^32^. Previous ssNMR data revealed that this μs dynamics is present in one of ubiquitin’s crystal forms.^33,34^

In addition to this internal motional process, we have recently provided evidence that ubiquitin molecules undergo overall “rocking” motion in the crystal.^35^ This motion was detected by a comparative ssNMR R_1ϱ_ relaxation measurement of different crystal forms, which revealed that in one of the crystal forms, termed cubic-PEG-ub, elevated relaxation rates throughout the molecule can be explained by an overall reorientational motion with an amplitude of several degrees. The presence of this motion was also confirmed by MD simulations. It was furthermore shown that the observed rocking has direct implications for X-ray crystallography: the crystal form that features rocking motion gives rise to low-resolution diffraction data and high Wilson B-factors in crystallographic experiments at 100 K. However, our previous study did not provide a precise estimate of the rocking motion time scale, and only a range from ∼100 ns to 100 μs was indicated.^35^

Here we investigate in parallel two different crystal forms, thus avoiding any bias that would arise from comparing dynamics measured by different methods (i.e., solution and solid-state methods). We use Bloch-McConnell-type R_1ϱ_ relaxation dispersion experiments, which were recently introduced to solid-state NMR^34^, to probe on a per-residue basis the dynamics-induced modulation of the chemical shift of ^15^N nuclei. In addition, we use R_1ϱ_ experiments at higher radio-frequency field strengths, near the rotary-resonance condition 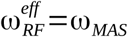,^34,36,37^which we refer to here as NEar-Rotary-resonance Relaxation Dispersion (NERRD) experiment. This experiment is sensitive to the angular fluctuations of the H-N bond vectors. A systematic analysis of experimental NERRD data against numerical simulations provides information about the angular motional amplitude of μs motions.

The combined experimental data establish that crystal contacts indeed alter the kinetics (exchange rates) and thermodynamics (relative populations of states) of μs motions. We find that the relative populations of the exchanging states, βI and βII, can be inverted in going from one crystal to another, and that in both analyzed crystal forms the exchange rates are slowed down significantly with respect to solution. Multi-μs long MD trajectories of WT and mutant proteins confirm these experimental findings, and identify the molecular origins of these altered dynamics. Using NERRD data, we establish that overall rocking occurs on the tens-of-microseconds time scale, with an amplitude of about 3-5 degrees. The crystal form that is amenable to rocking also features extensive βI/βII exchange. Furthermore, both rocking and exchange dynamics occur on a similar time scale, suggesting some degree of cooperativity between local and overall motions. Although our attempts to capture this effect by means of the rationally designed mutations proved unsuccessful, we believe that this scenario remains a possibility in this and other protein crystals. Our results highlight the importance of crystal packing effects in the context of crystallography-based investigation of protein dynamics.

## Results

## Theory

In order to probe μs dynamics we employ ssNMR ^15^N R_1ϱ_ relaxation dispersion (RD) experiments. RD experiments measure the transverse magnetization decay rate constant, R _1ϱ_, under an applied radio-frequency field, for each of the ^15^N backbone spins across the protein, and investigate how this decay rate constant depends on the field-strength, ν_RF_, of the applied RF field. In MAS ssNMR, two different physical mechanisms can give rise to (non-flat) RD profiles:

i. If the exchanging states differ in their local electronic environment around a given ^15^N nucleus, and if the exchange process occurs on a μs-ms time scale, the corresponding chemical-shift modulation contributes to the R_1ϱ_ decay rate constant. This enhancement of R _1ϱ_ can be scaled down by applying a sufficiently strong RF field, i.e. R_1ϱ_ decreases with increasing ν_RF_. This effect, known as “Bloch-McConnell RD” in solution-state NMR^38^, depends on the chemical shift difference between the exchanging sites, as well as the time scale of the exchange, the relative populations of the exchanging states and the amplitude of the RF field.
ii. Fluctuation of the dipolar and CSA interactions relative to the reference frame of the crystallite also induces enhanced R_1ϱ_ relaxation, in addition to the above effect that fluctuation of the isotropic chemical shift has.^34,37^ If the relevant motions are in the μs-ms range then this effect can be scaled down not only by application of the RF field, but also by magic angle sample spinning. Mathematically this is expressed by the formula derived by Kurbanov *et al^39^* which (in the case of sufficiently strong RF field surpassing the RF carrier offset 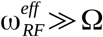 can be rendered in the following familiar form:

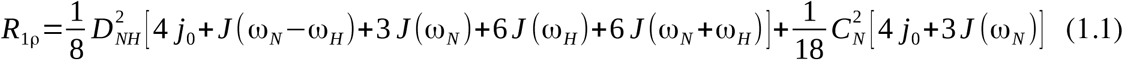

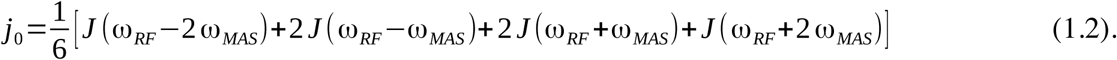

Here 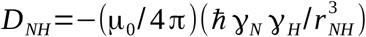 is the dipolar coup[ling strength, 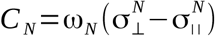 is the anisotropy of the chemical-shift tensor, ω*_RF_* =2 π ν*_RF_* is the spin-lock field strength, and ω*_MAS_*=2πν*_MAS_*. The explicit form of spectral densities *J*(ω) depends on the adopted model of motion; the empirical Lipari-Szabo-style expression is 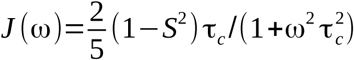.

Of particular importance in Eq. (1) are spectral densities *J* (ω_*RF*_±*n*ω_*MAS*_)(*n*=1,2). The meaning of these terms is as follows: if the motional process is sufficiently slow, ω*_RF_* τ*_c_*≥1 or ω*_MAS_* τ*_c_*≥1, then relaxation can be partially suppressed due to refocusing of the dipolar interaction by RF field or MAS, respectively. However, it may also happen that refocusing due to RF field and due to MAS in part cancel each other out (reminiscent of dipolar recoupling). This phenomenon occurs near rotary-resonance conditions, ω*_RF_* =*n* ω*_MAS_* or, equivalently ν*_RF_* =*n* ν*_MAS_* (*n* = 1, 2), and is accompanied by increase in the *R*_1ρ_ decay rate constant. Such behavior (illustrated by means of numerical simulations, Supplementary Figure 1) is the basis of NEar-Rotary-resonance Relaxation Dispersion (NERRD) experiment.

The observation of an increase of R_1ϱ_ when approaching the rotary-resonance condition is an unambiguous signature of μs motions. Assuming that dynamics can be modeled as an exchange process, the exact shape of such NERRD profiles depends on the time scale of motion, the relative populations of the involved states, as well as the difference in orientation of the NH bond vector between the exchanging states.^36,37^ NERRD thus provides insight into the angular amplitude of the motion, and complements Bloch-McConnell RD data, which probe motions by their isotropic chemical-shift modulation. Importantly, in solution-state NMR this type of information is unavailable, because the rapid (ns) molecular tumbling averages the anisotropic interactions such that they are not informative of slow dynamics.

Both of the above mechanisms, (i) and (ii), simultaneously lead to RF-field dependent R _1ϱ_ decay rate profiles whenever μs-ms motions are present. However, experimental parameters can be chosen such as to separate the two effects. Provided that a fairly large sample-spinning frequency *ν_MAS_* is used, the R1ρ RD profiles in the low RF-field range, typically ν_*RF*_≤10*kHz*, are dominated by isotropic chemical-shift mechanism. For simplicity, we employ the term “Bloch-McConnell relaxation dispersion” (BMCRD) for this situation. On the other hand, in the vicinity of the rotary-resonance conditions, which in our NERRD experiments is at ν_*RF*_=ν*_MAS_* =44 kHz and thus well separated from the BMCRD regime, it is the angular fluctuation of the NH bond that dominates the RD profiles.^34,37^ For simplicity, we will reserve the term NERRD for the situation where relaxation due to modulation of isotropic chemical shift is fully suppressed by the strong RF field and the near-resonance dispersion effect arises solely from the anisotropic interactions.

We note, however, that the two described relaxation dispersion phenomena (BMCRD, NERRD) are not fundamentally different but rather reflect two mechanisms of line broadening induced by μs motion. The two respective datasets can be fitted jointly or independently, and such fits can in ideal cases provide exchange rates, population levels of involved states, chemical-shift differences and jump angles. In systems where the exchange process is fast (< ∼100 μs), which is the case here, it is not possible to disentangle populations and chemical-shift changes, and only their product can be obtained; likewise, populations and jump angles also become entangled (see further discussion below and Supplementary Fig. 4).

## Crystal contacts slow down type-I/type-II β-turn exchange

In this study we investigate dynamics in two different crystal forms of ubiquitin which are obtained in the presence of two different precipitation agents (MPD and PEG, see Methods) and are referred to as MPD-ub and cubic-PEG-ub, respectively. X-ray diffraction based crystal structures of these different forms have been previously solved at cryo-temperatures (100 K). ^35,40,41^ While these X-ray data reveal that ubiquitin adopts essentially the same overall structure (backbone RMSD of 0.5 Å), they differ in the conformation of the loop region encompassing residues E51 to R54: in MPD-ub this region forms a so-called type-II β-turn, while it is in βI conformation in cubic-PEG-ub (Figure 1a). Note that the predominant state in solution is βI ^35,40-42^ with an estimated >95% population, in exchange with an alternate state which is believed to be βII.^30^

**Figure 1:**
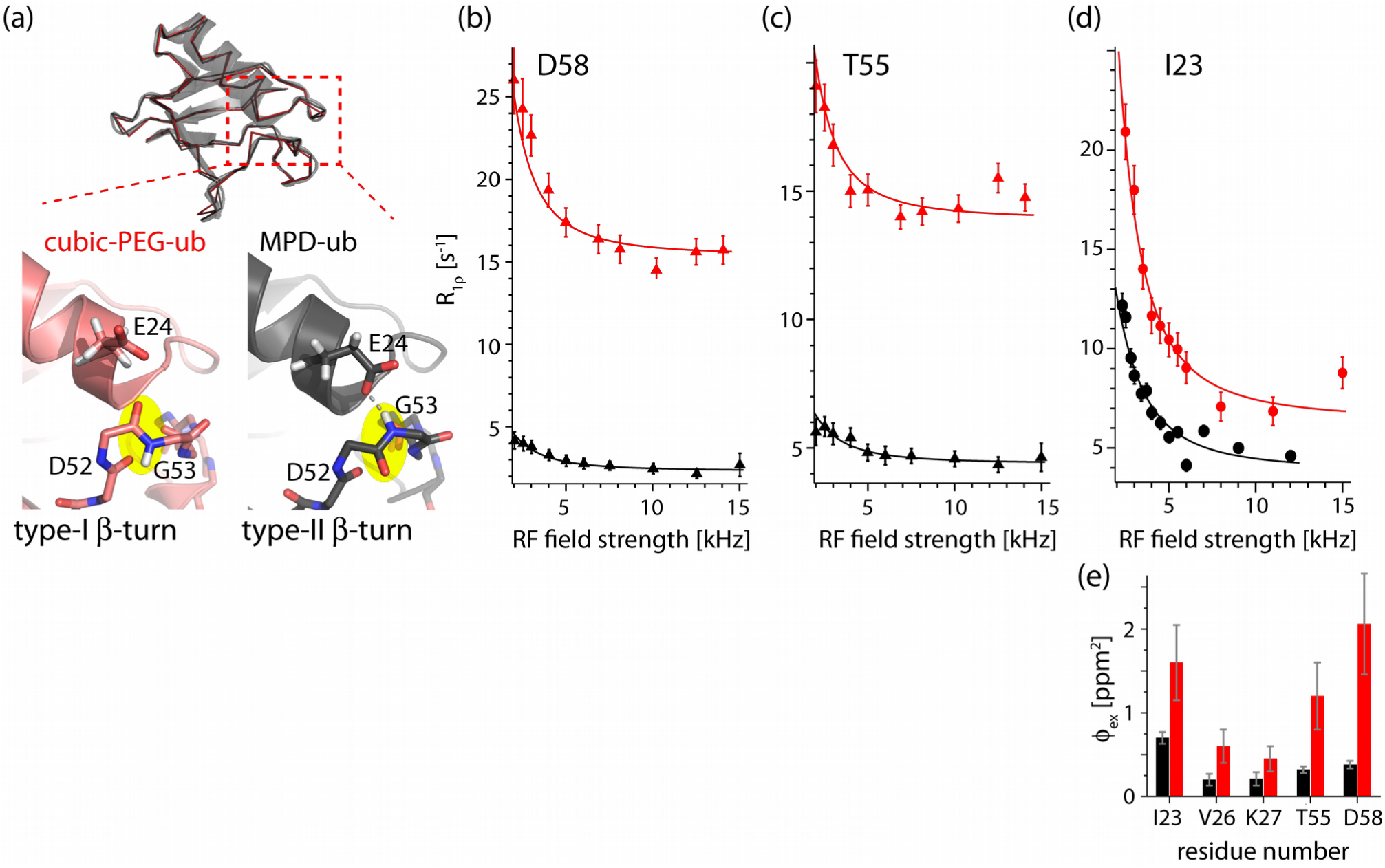
Evidence for β-turn exchange in MPD-ub (black) and cubic-PEG-ub (red) crystals. (a) Structures of ubiquitin in the two crystal forms. Top: Superposition of the main chains in the two crystals, shown as Cα backbone traces; for cubic-PEG-ub only one of the two non-equivalent chains is shown. For clarity the semi-transparent cartoon representation is shown only for MPD-ub. Bottom: zoom into the β-turn region. The peptide bond undergoing a flip is highlighted in yellow. (b-d) BMCRD profiles of ^15^N sites in the two ubiquitin crystal forms, obtained at a sample temperature of 300 K. Data shown here for D58 and T55 were obtained at B_0_ field strengths corresponding to ^1^H Larmor frequencies of 600 MHz, and those of I23 at 950 MHz, respectively. Error bars of R_1ρ_values were obtained by standard Monte-Carlo analyses, described in the Methods section. Solid lines correspond to a global two-site exchange model fit of data from residues I23, V26, K27, T55, D58 within each of the crystal forms (solid lines).The fits were performed separately for each of the two crystal forms using in each case the same cluster of the aforementioned five residues. Note that the resonances of E24 and N25 are not visible, presumably as a consequence of the conformational exchange. The comprehensive set of BMCRD data and fitting results can be found in Supplementary Figures 2, 3, and 5. (*e*) *ϕ_ex_* values, 2 ϕ*_ex_* = *p_A_ p_B_* (2 π Δ δ) in cubic-PEG-ub (red) and MPD-ub (black), obtained from the joint fit of residues I23, V26, K27, T55 and D58, exemplified by curves in panels b-d. Here *p_A_* and *p_B_* are the populations of the two conformers, while Δδ is the chemical shift difference expressed in the units of ppm. Fitted exchange time constants are reported in Table 1.

**Table 1.**
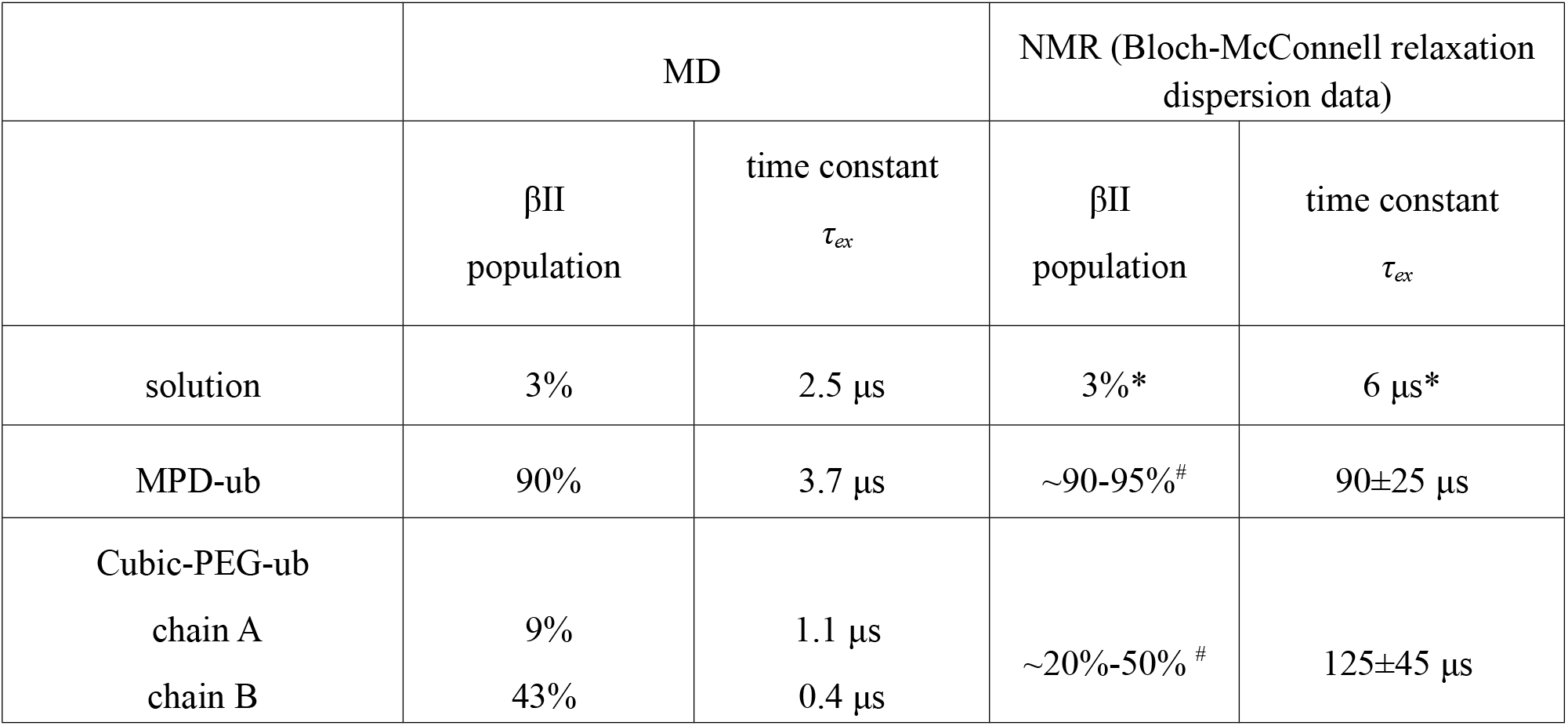
Populations and time scales of conformational exchange dynamics in the region encompassing residues L50-L56 and the adjacent N-terminal part of the α-helix. The population of the βI state is p_βI_ = 100%-p_βII_.

Figure 1b-d shows Bloch-McConnell RD data for three representative amide sites in MPD-ub and cubic-PEG-ub. The non-flat RD profiles for residues located in the β-turn region and the adjacent N-terminal part of the α-helix unambiguously show that this part of the molecule undergoes μs dynamics in both crystals. A joint fit of a two-state exchange model to data from the five residues located in this region is shown as solid lines (see Methods). The use of a two-state exchange model appears justified in the light of the two distinct states observed in different crystal structures and MD simulations (ref. ^32^ and MD data below). The exchange time constant obtained from these fits is τ_ex_=90 ±25 μs for MPD-ub and 125±45 μs for cubic-PEG-ub. We want to stress the fact that the absolute “plateau” levels of these curves are significantly higher in cubic-PEG-ub than in MPD-ub, throughout the molecule, as can be readily seen in the three examples in Figure 1b-d. While the plateau is not relevant for fits of BMCRD data, this finding of elevated R _1ρ_ rates points to the presence of overall angular motion of ubiquitin molecules in cubic-PEG-ub, as we will outline below.

In order to gain insight into the angular motions of the NH bond vectors, we measured NERRD profiles. Figure 2 shows the largest NERRD effects observed in MPD-ub, which are found for the amide sites of D52 and R54, which are adjacent to the peptide plane D52/G53 that undergoes a ca. 180° flip when exchanging between βI and βII conformations. (The amide signal of G53 is not resolved in the spectra of MPD-ub.) Other residues do not show significantly rising NERRD profiles, as exemplified by the data of three residues depicted in black in Figure 3. An independent experiment at a different MAS frequency (20 kHz, vs. 44 kHz in the current experiment) also exhibited the largest NERRD effect for residues D52 and R54 in MPD-ub ^35^, whereas all other residues showed either small NERRD profiles (residues I23, E51, T55) or no significant NERRD effect, mirroring the present data.

**Figure 2.**
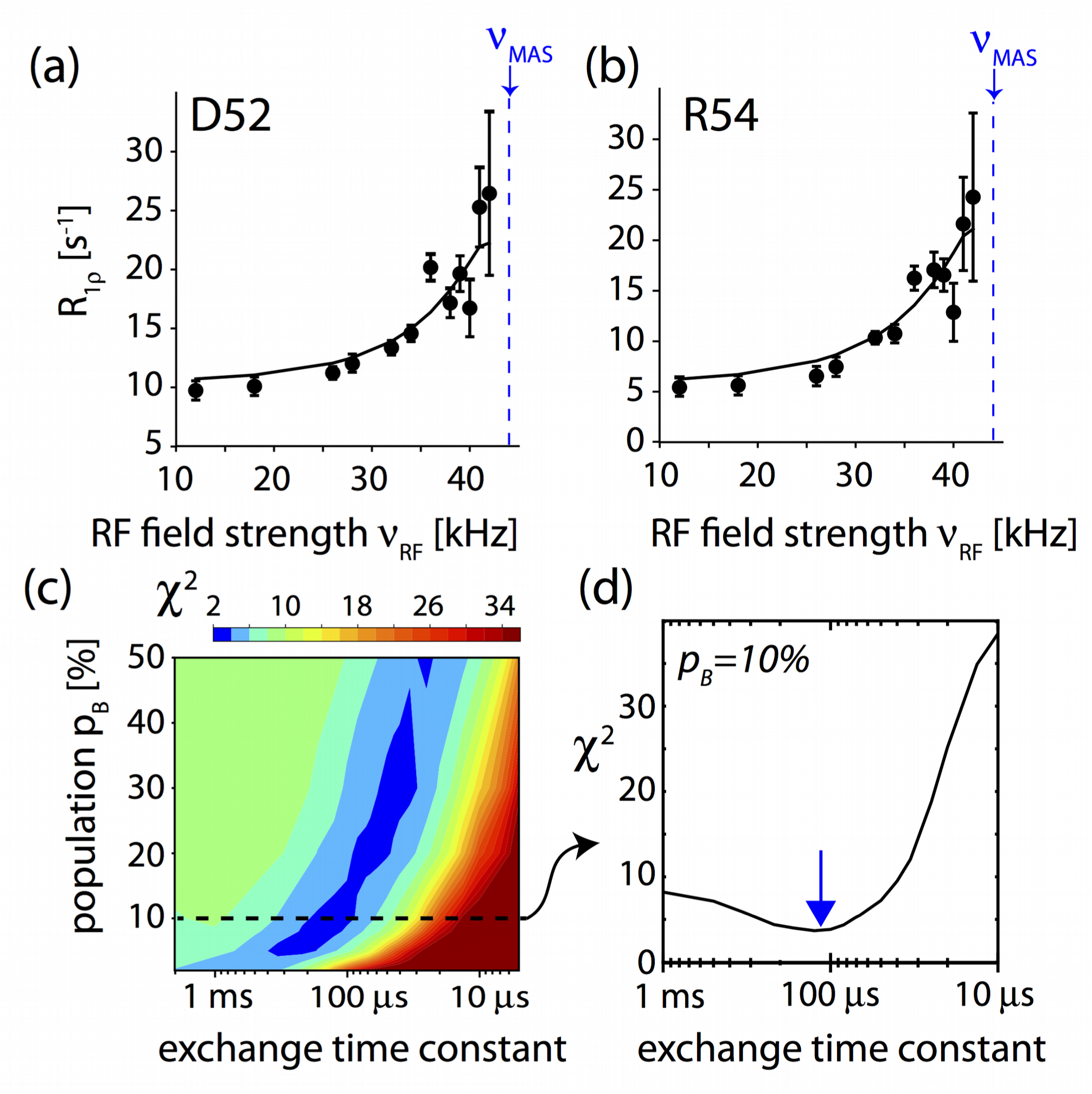
Angular motion in the β-turn of MPD-ub crystals probed by NERRD NMR. (a,b) NERRD dispersion data of the two amide sites in MPD-ub that are adjacent to the flipping peptide plane D52/G53, recorded at 44.053 kHz MAS frequency. Error bars of R_1ϱ_ values were obtained by standard Monte-Carlo analyses, described in the Methods section. Solid lines show fit curves from a two-site jump model that assumes that the minor conformation has a population of p_B_=10%. (c) Chi-square surface of a grid-search of simulated NERRD curves against the experimental data in panels (a,b). Hereby, a common minor-state population p_B_ and exchange correlation time τ_ex_=1/ (k_AB_+k_BA_) were assumed for the two residues, and the jump angle was separately fitted for each residue. More details of this NERRD fit are reported in Supplementary Fig. 4. (d) Cross-section across the chi-square surface along the τ_ex_ dimension corresponding to a minor-state population of 10%. This minor-state population level is suggested by analyses of BMCRD data, as explained in the text. The best-fit exchange time constant with this assumption is τ_*ex*_ ∼ 100 *μ*s.

**Figure 3.**
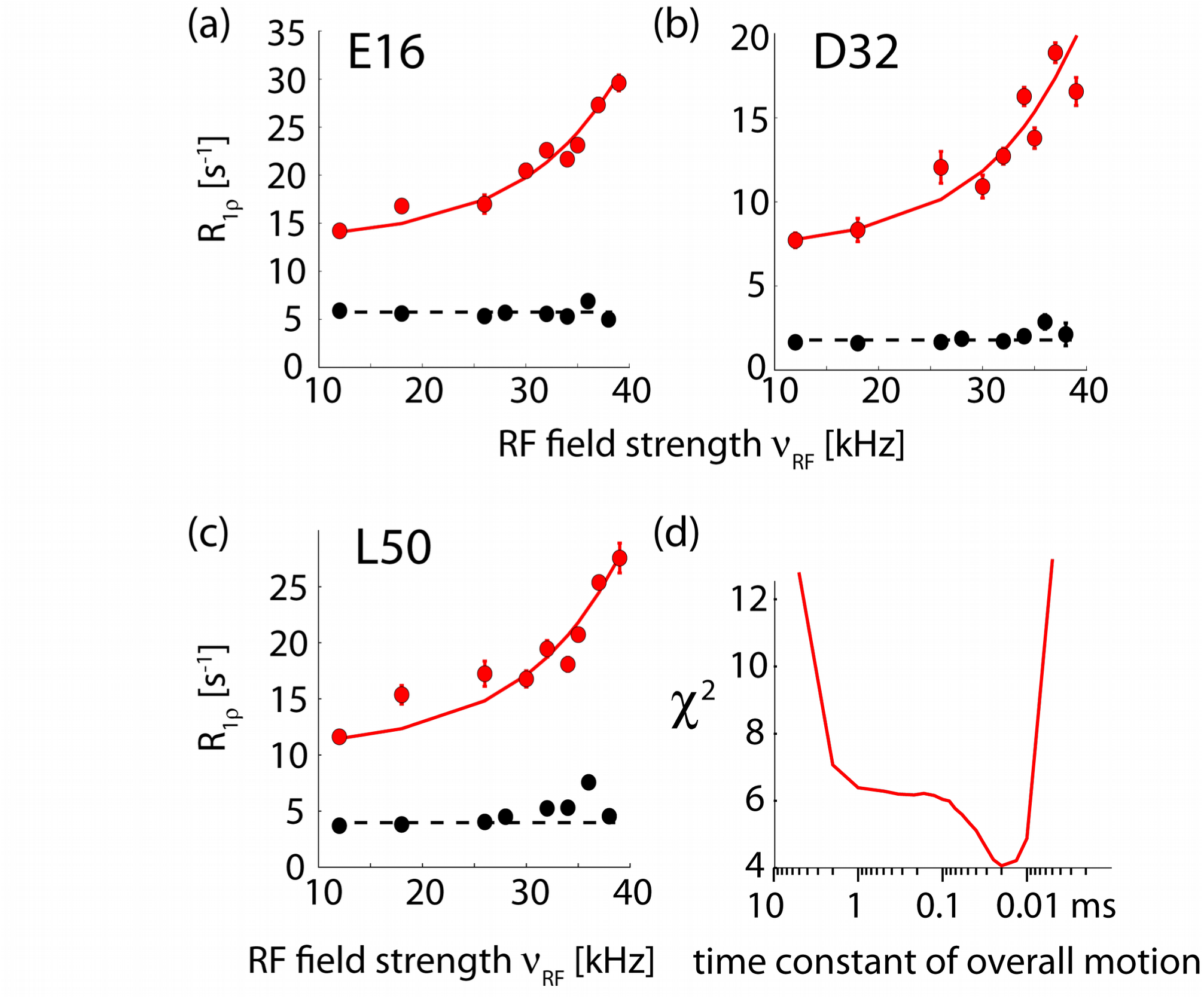
Evidence for rocking motion in cubic-PEG-ub crystals from NERRD NMR measurements. (a-c) NERRD data of representative amide sites outside the dynamic β-turn region for MPD-ub (black) and cubic-PEG-ub (red). Error bars were obtained from Monte Carlo analysis, described in the Methods section. The solid red lines correspond to a two-state fit of data from 22 amide sites with a common exchange time constant and per-residue-adjustable motional amplitude (all shown in Supplementary Fig. 6). (d) χ^2^ value of this fit as a function of the jump time constant. In fitting these NERRD data we have assumed that protein rocking can be approximated as two-site jump process, which is a crude but justifiable model.^47^

To quantitatively rationalize these data we used numerical spin-dynamics simulations to obtain theoretical NERRD profiles over a grid of exchange parameters, *p_B_* and *τ_ex_*, and fitted the experimental data against this grid of simulated curves. Solid lines in Figure 2a,b show the best-fit curves for such a joint fit of data from D52 and R54, assuming that they have one common excited-state population and exchange time constant. Figure 2c shows the corresponding plot of the χ^2^ surface. Although the population levels obtained from these fits have significant uncertainty, the time scale converges to ∼50-200 μs. This is in excellent agreement with the independent Bloch-McConnell RD data, as illustrated in Figure 1b-d. In addition to confirming the time scale of the process, NERRD data provide information about the amplitude of the NH motion of D52 and R54, which is found to be on the order of 7 to 11 degrees, depending on the assumed populations (see Supplementary Fig. 4). Interestingly, this angle is similar to the difference between the orientations that these two bonds assume in βI and βII states, which is 12 and 14° for the two respective residues (comparing PDB entries 1UBQ and 3ONS).

Taken together, both Bloch-McConnell RD and NERRD data show that the β-turn dynamics in MPD-ub occurs on a time scale of ca. 100 μs; this exchange process causes chemical shift modulation in the β-turn region (Figure 1) and angular jumps of the amides of D52 and R54, as revealed by NERRD (Figure 2).

It is interesting to note that the two crystals, which have very different packing environment around the exchanging region, have similar exchange kinetics, but that these kinetics are rather different to the case of solution, where the exchange process has a time constant of τ _ex_=6 ± 2 μs at the same temperature, i.e. is about 15 times faster.^30^ The significant slowdown of the exchange process in the crystals is not due to the increased viscosity induced by the precipitation agent: it has been shown^33^ that even in the presence of MPD concentrations as high as 45% in solution all CPMG RD profiles were flat, i.e. the underlying process was too fast to induce RD, whereas in MPD-ub crystals very large CPMG RDs were detected, thus confirming that MPD is not the principal cause for slowing of the exchange dynamics.^33^ As noted above, the “ground state” is different between the two crystals, being βI in cubic-PEG-ub (as well as in solution), and βII in MPD-ub. Our findings show that the nature of the ground state does not dictate the kinetics, because in solution and cubic-PEG-ub, which have both βI as major state, the kinetics are very different.

## βI/βII populations differ in the different crystals

Inspection of the BMCRD data in Figure 1 reveals a noticeable difference between the two crystals: the amplitude of these RD profiles for residues in cubic-PEG-ub is considerably larger than those of their counterparts in MPD-ub, on average by a factor 3.3±1.0 with regard to the values of ϕ*_ex_* in Figure 1e.

While precise determination of populations is beyond reach in the case of fast exchange, semi-quantitative estimates are nevertheless possible. ϕ_ex_ values in MPD-ub are consistent with *p*_β_*_I_* ≈5−10% and Δδ on the order of a fraction of ppm, as can be expected for the analyzed residues that are located at the periphery of the β-turn. This *p*_β_*_I_* estimate is supported by our MD simulations (see below). It is further reasonable to assume that Δδ values are approximately the same for the MPD-ub and cubic-PEG-ub experiencing the same exchange process (interconversion between βI and βII). If we make this assumption, then based on ϕ_ex_ results we can estimate that the populations of minor species in the cubic-PEG-ub crystal is *p*_β_ *_II_* ≈20−50%. Note that the population of minor species cannot exceed 50%, which provides in this context a useful boundary. Supplementary Table 1 provides some example calculations relating the minor-state populations in the two crystals.

MD simulations using explicit crystal lattices have previously been shown to successfully reproduce experimental ssNMR and XRD data, and they may contribute valuable information about mechanistic details of motion that are difficult to access experimentally. ^6,35,43-45^ We simulated an explicit arrangement of ubiquitin molecules according to the crystal lattices of MPD-ub and cubic-PEG-ub, built of 24 and 48 ubiquitin molecules, respectively, in the presence of interstitial water over a time interval of 2 μs. The explicit simulation of a large number of ubiquitin molecules improves the sampling statistics considerably. In addition, we performed a 10 μs simulation of ubiquitin in solution, which we find to be in excellent agreement with a recent 1 ms trajectory. ^32^ In all these trajectories we observed multiple βI ↔ βII transitions, enabling us to extract the relative populations of the two conformations, which are reported in Table 1. The minor-state populations are qualitatively in good agreement with the estimates derived from BMCRD data, i.e. simulated MPD-ub has significantly lower minor-state population than cubic-PEG-ub (chain B), mirroring the lower ϕ_ex_ values. It should be noted that in the ssNMR spectra of cubic-PEG-ub the signals from the two inequivalent chains, A and B, overlap in most cases, making it impossible to collect two separate datasets. The experimental data, therefore, reflect an effective average of the dynamics in the two chains. With this caveat the *p*_β_ *_II_* populations found in the MD trajectory of cubic-PEG-ub are consistent with the experimental observations.

The time scale of the exchange process in the MD simulations tends to be faster than observed experimentally (see Table 1), suggesting that the force field employed in this study does not accurately reflect the transition-state free-energy barrier. Similar findings have been reported before.^46^ This observation is perhaps not surprising, given the limited accuracy of MD models (note that force fields have been parameterized primarily against experimental equilibrium data, and not against kinetic data). From the perspective of this study the elevated exchange rates are to some degree advantageous since this improves the statistical sampling of the βI/βII equilibrium.

Thus, NMR and MD techniques independently indicate that the relative populations of βI and βII conformers differ considerably in different crystals and in solution. The βI:βII populations span the wide range from 10:90 (MPD-ub) to 57:43 / 91:9 (cubic-PEG-ub) and 97:3 (solution).

## Molecules in cubic-PEG-ub undergo tens-of-µs rocking motion

We also recorded NERRD experiments on cubic-PEG-ub crystals, in order to obtain additional insight into μs dynamics in this crystal form. It shall be reminded that in MPD-ub only few residues (in particular D52 and R54; see Fig. 2a,b) show increased R _1ϱ_ close to the rotary resonance condition (i.e. non-flat NERRD profiles), and that the time scale obtained from a fit of these data is in excellent agreement with Bloch-McConnell RD data. In contrast to this finding, essentially all residues in cubic-PEG-ub exhibit non-flat NERRD profiles, as shown in Figure 3 and Supplementary Fig. 6, and the R_1ϱ_ values are generally higher than those observed in MPD-ub. Given that this trend is observed for all residues, we ascribe the underlying motion to a global process involving the entire molecule, such as an overall “rocking” motion of the ubiquitin molecules in the crystal. Overall rocking motion is visible not only in R _1ϱ_ measurements, but is also expected to contribute to line widths in ssNMR experiments. This expectation is indeed confirmed by ^1^H and ^15^N coherences life times, which differ markedly between the two crystal forms studied here (Supplementary Fig. 7). We have proposed the existence of such overall motions in cubic-PEG-ub recently, based on the observation of elevated R_1ϱ_ rate constants in cubic-PEG-ub compared to MPD-ub,^35^ although the time scale could be estimated only very roughly (ca. 100 ns to 100 μs).

In order to obtain quantitative insight into rocking motion, we have performed a common fit of a dynamic model with a global time scale to NERRD data from 22 well-resolved residues in cubic-PEG-ub, excluding residues in the β-turn region, since for those residues one expects superposition of local and global motion. The best-fit time scale is of the order of tens of microseconds (Figure 3d). Residue-wise motional amplitudes, expressed in terms of a two-site jump model, are in the range 3-5° (Supplementary Fig. 6). Such a relatively small amplitude is expected for an overall-rocking motion within the confines of the lattice; MD simulations revealed an amplitude of overall motion in cubic-PEG-ub of ∼6-8°, although this amplitude is likely exaggerated due to simultaneous small drift (“melting”) of the crystal lattice.^35^

It is interesting to note that the tens-of-microseconds time scale of overall motion, found experimentally, is similar to the time scale of the β-turn conformational exchange in cubic-PEG-ub (125±45 μs). This similarity may be fortuitous, but it might also point to some correlation between the two processes, i.e. the local and the overall motion. Such possibility is discussed in what follows.

## Intermolecular contacts that alter conformational equilibria

As noted above, the βI conformation in ubiquitin represents a “default” state, which is found in solution and in the majority of crystal structures. At the same time the βII conformation has also been observed in a number of crystals, such as MPD-ub ^41^ and complexes of ubiquitin (see refs. in Sidhu *et al^48^*). This leads us to conclude that the crystalline environment can act as a switch converting the protein from βI to βII. Our MD simulations of ubiquitin crystals are well suited to identify intermolecular interactions that are responsible for this transformation.

In our initial analysis of the MPD-ub trajectory we have identified two such putative intermolecular contacts. One is the four-way junction involving a pair of D52 side chains and a pair of K63 side chains, all of which belong to different ubiquitin molecules but come together in the crystal lattice (Figure 4a). The other is the intermolecular hydrogen bond between the (neutral) carboxyl groups of E64 and E24, which appears to stabilize the coupling between E24 and G53 (Figure 4b). According to the MD simulation data, both of these interactions show clear preference for the βII over the βI form.

**Figure 4.**
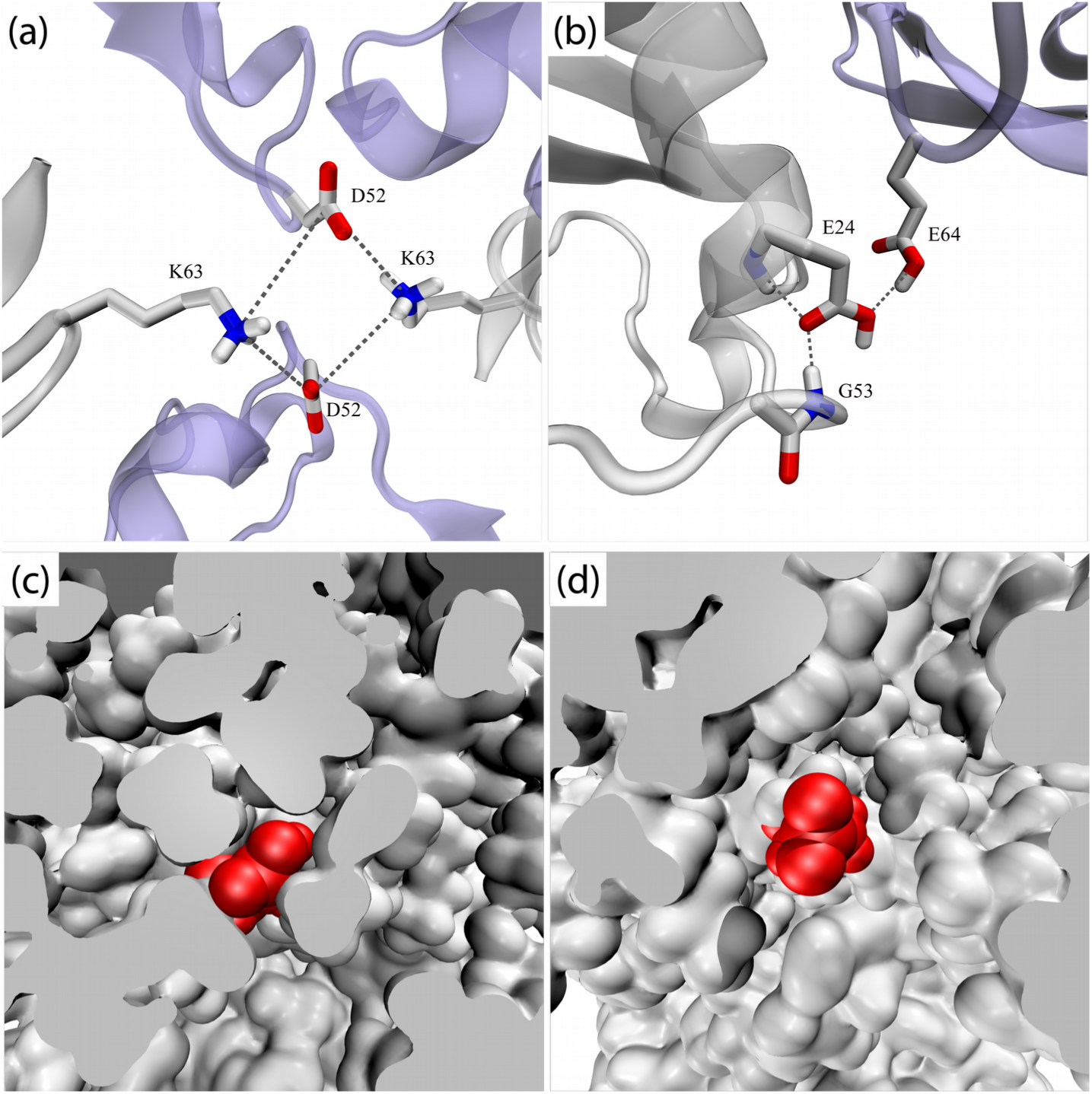
Inter-and intra-molecular interactions that affect β-turn motion. (a), (b) Intermolecular interactions involving the β-turn region in MPD-ub. Neighboring molecules are shown in light blue. Simulations of ubiquitin mutants suggest that these interactions are not responsible for causing the preference for βII, see text. (c), (d) The surrounding of E24 side chain in different crystal forms of ubiquitin. (c) A rare frame from the MPD-ub trajectory where E24 side chain (painted red) is projected toward solvent. (d) A typical frame from cubic-PEG-ub trajectory where E24 from ubiquitin molecule chain A is immersed into solvent.

However, additional simulations convinced us that this is the case of correlation rather than causation. Specifically, we have recorded two separate 2-μs-long trajectories of MPD-ub crystals, one of which carried K63A mutation and the other E64A mutation. As it turned out, these trajectories produced exactly the same proportion of βI to βII species as the wild-type crystal simulation. This led us to conclude that the intermolecular contacts involving K63 and E64 side chains are not the key to the β turn conformation.

At the same time we have found that an E24A mutation produced a complete reversal of βII to βI ratio compared to the wild-type simulation, from 90%:10% to 9%:91%. In other words, E24A mutation has restored the “default” state of the system, which is dominated by βI. Apparently it is the hydrogen bond between the carboxyl group of E24 and the backbone amide group in G53 which is directly responsible for prevalence of the βII form in the wild-type MPD-ub crystal. But this hydrogen bond is an *intra*molecular hydrogen bond – so how is it related to crystal packing?

Inspection of the trajectories of ubiquitin in solution and in the cubic-PEG-ub crystal revealed that in these simulations the side chain of E24 is mostly immersed in solvent. In solution, E24 side chain is solvated 65% of the entire simulation time. In cubic-PEG-ub, it is solvated 62% of the time for chain A and 52% of the time for chain B. In contrast, it is solvated at the level of only 11% in the MPD-ub simulation (see Table 2).

**Table 2.**
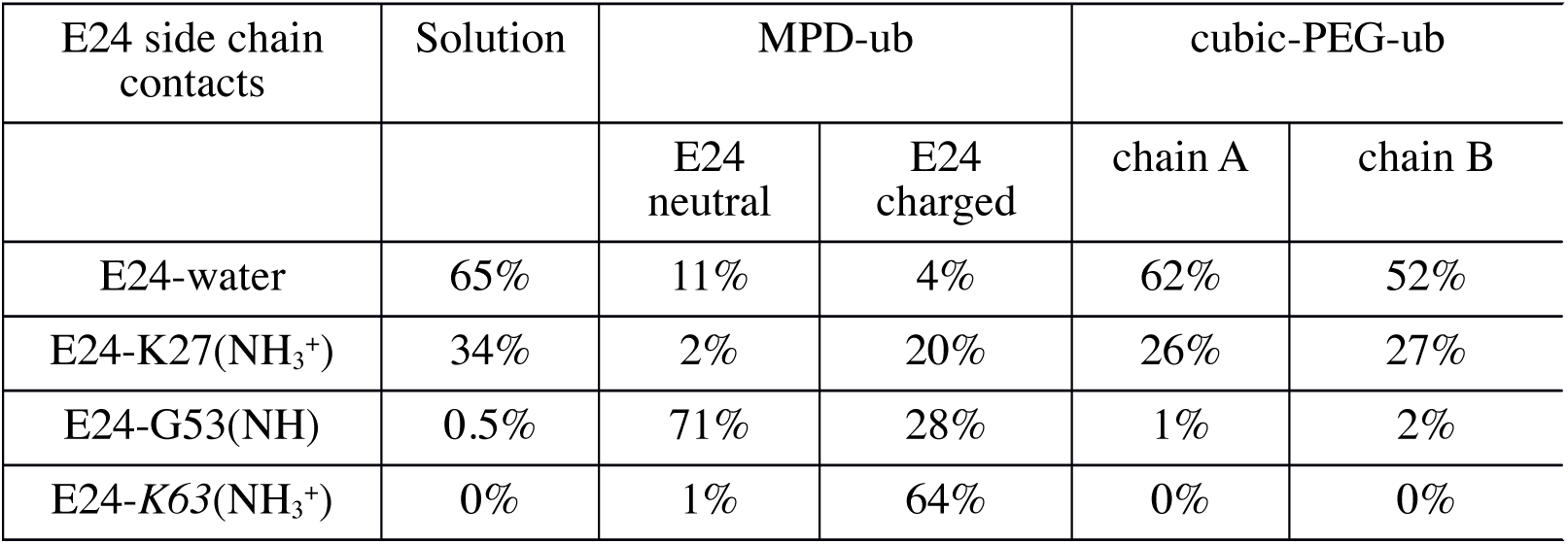
Important interactions involving E24 side-chain carboxylic group as characterized by their presence in the MD simulations. The contacts involving K27 and G53 are intramolecular, the contact involving K63 is intermolecular. The numbers in each column do not always add up to 100% because E24 also forms certain other contacts (both inter-and intramolecular) or, otherwise, some of the listed interactions can occur simultaneously.

| E24 side chain contacts | Solution | MPD-ub |  | cubic-PEG-ub |  |
| --- | --- | --- | --- | --- | --- |
|  |  | E24 neutral | E24 charged | chain A | chain B |
| E24-water | 65% | 11% | 4% | 62% | 52% |
| E24-K27(NH<sub>3</sub><sup>+</sup>) | 34% | 2% | 20% | 26% | 27% |
| E24-G53(NH) | 0.5% | 71% | 28% | 1% | 2% |
| E24-K63(NH<sub>3</sub><sup>+</sup>) | 0% | 1% | 64% | 0% | 0% |

As it turns out, the packing of ubiquitin molecules in the MPD-ub crystal lattice is such that E24 side chain does not have the space to extend outwards. This is illustrated in Figures 4c,d, which compare the volume of space available to E24 side chain in MPD-ub and in cubic-PEG-ub crystals. In particular, the intrinsically favorable rotameric state (χ_1_=180°, χ_2_=180°) which is prominent in other simulations, is completely obstructed in MPD-ub. We conclude that typically the E24 side chain enjoys the entropic benefit of being immersed in solvent, but in MPD-ub it is denied this opportunity and therefore opts for the hydrogen bond with G53.

The propensity of the carboxylic group in E24 to form a hydrogen bond with the G53 amide group is also influenced by its protonation state. The MPD-ub crystal has been obtained from a mother liquor at pH 4.2. Under these conditions the carboxylic group of E24, as positioned in the crystallographic structure, is expected to be mostly protonated ^49^. To elucidate the role of this factor, we have altered E24’s status from neutral to anionic and then recorded an additional 2-μs-long simulation of the MPD-ub crystal. As it turns out, in this trajectory E24 is partially recruited into both intra-and inter-molecular salt bridges, see Table 2. This happens at the expense of the E24-G53 hydrogen bond, which drops to the level of 28%. Accordingly, the ratio of βII to βI in this trajectory changes from 90%:10% to 71%:27%. This finding supports the hypothesis that the protonation state of E24 also plays a significant role in the stabilization of the βII species in MPD-ub crystal.

### Potential coupling between rocking and βI/βII exchange

As discussed above, cubic-PEG-ub crystal is characterized by significant amount of rocking and, at the same time, extensive conformational exchange in the βI/βII region. Furthermore, the characteristic time constants of these two motional processes turn out to be similar. Is this a mere coincidence, or does it point toward a certain common underlying mechanism?

Inspection of the cubic-PEG-ub trajectory led us to identify one potential coupling mechanism. This mechanism involves an ion pair between K11 (chain A) and D52 (chain B). In the crystal structure these two side chains are positioned far apart (Figure 5a). However, during the MD simulation they frequently form a salt bridge (Figure 5b). The formation of this salt bridge is associated with (*i*) slight reorientation of protein molecules in the crystal lattice and (*ii*) βI → βII transition mediated by residue D52 in the β turn. Thus the reorientational dynamics of the protein (rocking motion) may be correlated with conformational exchange. This hypothetical scenario is illustrated in the Supplementary Movie 1.

**Figure 5.**
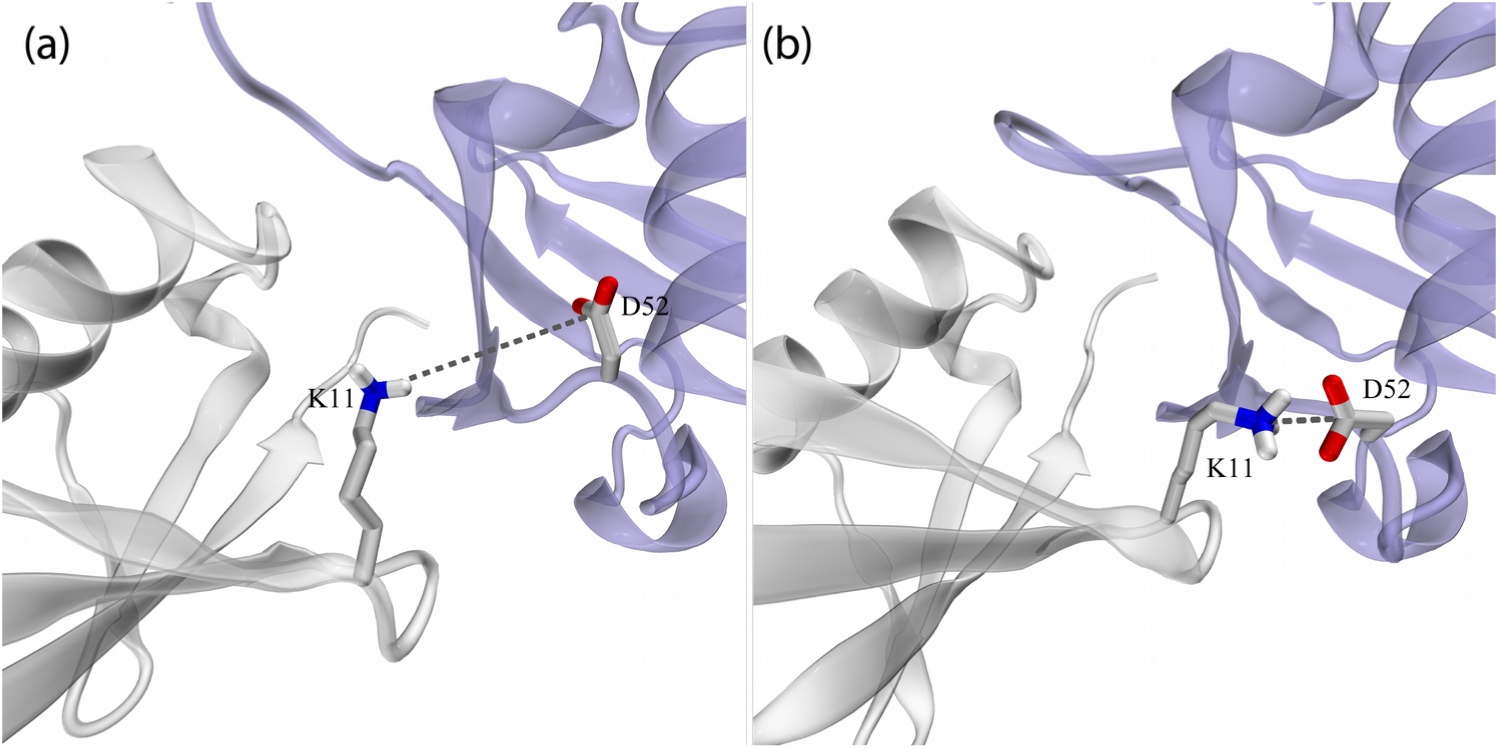
Dynamic ion pair K11_chainA_ – D52_chainB_ interaction in cubic-PEG-ub crystals. (a) Crystallographic structure of cubic-PEG-ub (PDB entry 3N3040) and (b) a selected frame from cubic-PEG-ub trajectory. The dynamic disorder in residue K11_chainA_ is reflected in its crystallographic temperature factors; this residue’s side chain has not been modeled in our recent structure 4XOL.

We have yet been able to obtain only partial confirmation for this scenario. The MD data indicate that K11_chainA_-D52_chainB_ salt bridge in the cubic-PEG-ub crystal favors βII conformation over βI (by a factor of 4.1 considering the trajectory at hand). In order to probe this effect experimentally, we have prepared cubic-PEG-ub crystals of K11A ubiquitin, and performed BMCRD experiments. We saw a small but detectable decrease in RD amplitudes for residues in the β-turn region in the K11A ubiquitin crystal compared to WT crystal (Supplementary Figure 10). This reduced RD effect may indeed point to a decreased population of minor βII state in K11A.

On the other hand, our attempts to detect the link between K11 _chainA_-D52_chainB_ and rocking motion proved to be unsuccessful. The MD simulation of K11A crystal failed to produce any significant evidence that on-off K11_chainA_-D52_chainB_ salt bridge drives rocking dynamics. In part, this can be attributed to poor convergence of the MD trajectories, which are affected by gradual “melting” of the crystal lattice (discussed by us previously;^35^ see Supplementary Figure 12). One should also bear in mind that ubiquitin molecules in the crystals are held together by a complex network of intermolecular interactions involving several dozen hydrogen bonds and salt bridges (see Supplementary Figures 8 and 9). In this situation the effect from the single point mutation, K11A, may be masked; moreover, the consequences of the mutation may be more complex and differ from what has been expected. The experimental NERRD data from the sample of K11A ubiquitin also could not detect the expected decrease in rocking within the experimental precision (Supplementary Figure 11).

Although in this specific case we could not confirm that conformational exchange is coupled to rocking, this hypothesis deserves further investigation. It is envisioned that local conformational rearrangements may lead to “repacking” of protein molecules in the crystal lattice, entailing small overall translation and reorientation (i.e. causing rocking motion). Conversely, rocking motion may lead to subtle changes in the pattern of crystal contacts, triggering local conformational transitions. In general, it should not be surprising that a molecule that is conformationally labile produces a poor crystal which suffers, *inter alia*, from rocking dynamics.

## Probing relaxation dispersion effects

MD simulations on a microsecond time scale can be used to model BMCRD effects. For this purpose MD coordinates are fed, frame by frame, into chemical shift prediction programs such as SHIFTX.^50^ In a situation when the calculated chemical shift δ(^15^N) shows a pattern of modulation on the μs time scale, one may expect to experimentally detect RD effects at the respective site.^51^ Ultimately, this approach makes it possible to uncover the dynamic mechanism behind the observable RD profiles.

We have used this method to probe the backbone ^15^N sites in MPD-ub and cubic-PEG-ub trajectories. In MPD-ub, relaxation dispersion effects have been predicted for sites in the β-turn experiencing βI ↔ βII exchange, as well as sites at the N-terminal end of the first α helix that are hydrogen-bonded to this β turn. One example of such behavior, residue R54 from MPD-ub, is illustrated in the Supplementary Movie 2, demonstrating that the βI ↔ βII transition is accompanied by a ^15^N chemical-shift change in R54.

Similarly, the analysis of the cubic-PEG-ub trajectory suggests that Bloch-McConnell dispersions can occur in the β turn and its coupled sites (particularly, in the chain B molecules). In addition, we have also found several sites where the origin of predicted RD effects is different. One example of such distinctive behavior is residue H68. Focusing on one specific ubiquitin molecule in cubic-PEG-ub trajectory, we observe H68 side chain engaged in hydrogen bond with Q40 side chain from a neighboring molecule (Figure 6a). Later during the simulation H68 side chain swings sideways and forms hydrogen bond with side chain E64 which belongs to another neighboring molecule (Figure 6b). After ca. 0.8 μs it makes a reverse transition, reestablishing hydrogen bond with Q40, and remains in this conformation until the end of the trajectory.

**Figure 6.**
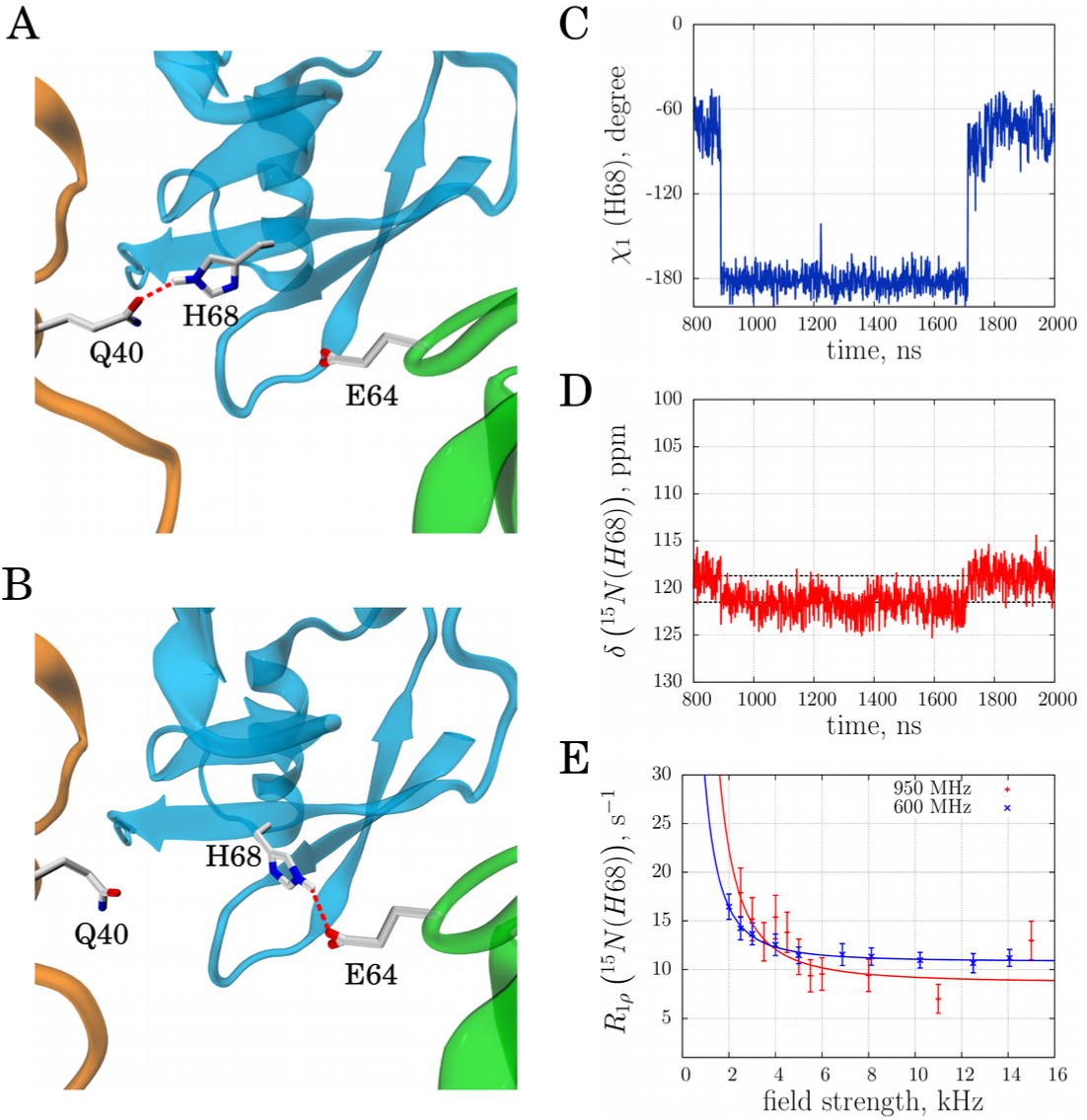
μs time scale dynamics of residue H68 (chain A) in cubic-PEG-ub crystals. (a,b) Intermolecular hydrogen bonds alternatively formed between H68 and Q40 or E64 as seen in the MD simulation of cubic-PEG-ub. (c,d) Time variation of side-chain torsional angle χ_1_(t) and main-chain ^15^N chemical shift δ(t) for residue H68 in the selected ubiquitin molecule from the simulated unit crystal cell. (e) Experimental relaxation dispersion data for residue H68 (chain unassigned) in the cubic-PEG-ub crystal; also shown are the results of fitting using fast exchange model (continuous lines). Error bars of R1ρ values were determined from Monte Carlo analysis. The motional time scale according to the experimental data, 56 μs, is one-to-two orders of magnitude slower than suggested by the MD simulations (the same is true for other manifestations of μs dynamics in ubiquitin crystals). The animated version of this graph is available as Supplementary Movie 3.

The rotameric transitions of H68 (from χ_1_=-60° to 180° and back) cause modulation of the backbone chemical shifts in this residue. For the ^15^N spin the amplitude of modulation is calculated to be 2.1 ppm (corresponding to the gap between the two dashed horizontal lines in Figure 6d). This is a sizable effect, which can be reliably reproduced by chemical shift prediction software. Therefore the computational analysis points toward possible RD effect at the amide site in residue H68. This prediction is borne out by the experimental data illustrated in Figure 6e.

Of interest, there is a theoretical possibility that conformational exchange in H68 is influenced by rocking motion (cf. previous section). Indeed, slight reorientation of the central protein molecule relative to its neighbors should favor either H68-Q40 or H68-E64 interaction depending on the direction of reorientational fluctuation (see Figures 6a,b). In this connection it is noteworthy that RD effects have been experimentally detected at this site in the cubic-PEG-ub crystal form which is affected by rocking, but not in the MPD-ub crystal which is immune to rocking (see Supplementary Fig. 3).

## Discussion

The advent of novel experimental and theoretical tools to decipher the heterogeneity of structures co-existing in crystals has sparked increased interest in molecular motions of crystalline proteins and the role of crystal packing in the context of dynamics. Here we have obtained a comprehensive portrayal of μs dynamics in different crystal forms of ubiquitin. Experimental ^15^N R_1ϱ_ relaxation data, specifically the combination of BMCRD and NERRD analyses, produced evidence of two major motional modes: conformational exchange in the β-turn formed by residues E51-R54 and overall rocking. The characteristic time scale of a local β-turn interconversion motion in the two investigated crystals is ca. 100 μs, more than one order of magnitude slower than in solution. Overall rocking motion entails small reorientational fluctuations of the protein molecules within the confines of the crystal lattice and affects molecules in one of the two crystal forms, cubic-PEG-ub, much more significantly than in the other, MPD-ub. The amplitude of the rocking motion in cubic-PEG-ub was determined to be ca. 4°, and the time constant was found to be on the scale of tens of μs. These results, as well as other findings reported in our study, show that μs dynamics is dependent on the environment.

MD simulations, including mutant simulations, can rationalize these effects. In particular, we can tentatively explain why MPD-ub prefers the type-II β-turn conformation. The acquired experimental evidence together with MD data hint at the possibility that internal conformational dynamics may be coupled to the rocking motion. Elucidating this connection in protein crystals is a daunting task. In particular, accurately modeling something as subtle and fickle as crystal contacts is a major challenge for the existing MD methodology. Nevertheless, we expect that future advances in the field of MD simulations should eventually make it possible. The identification of intermolecular interactions that control overall motion, either directly or through their involvement in internal conformational exchange, might open future possibilities for rational design of mutant proteins that could reduce the effect of rocking and thus improve diffraction quality.

It is known that internal dynamics undermine the quality of X-ray diffraction data. Crystallographers go a great length in order to reduce the level of dynamics in their samples (e.g. by engineering point mutations or by introducing ligands). It has not been fully appreciated, however, that internal conformational exchange may be connected to rocking motion, which can be even more detrimental for crystallographic analyses. We have shown here that both internal and overall motions can be probed by ssNMR spectroscopy and MD simulations. It is envisaged that both methods will be successfully exploited in future to either help in designing of stable crystals or, conversely, to assist in X-ray diffraction based studies of protein motions.

## Methods

## Sample preparation

Uniformly ^2^H,^15^N-labeled human ubiquitin (without any affinity tag) was over-expressed in *E. coli*, and purified using anion exchange and size exclusion chromatographies. In order to have a dilute ^1^H spin network and thus minimize the possible perturbing effect of remote ^1^H spins on the relaxation of amide ^15^N, the protein was dissolved in H _2_O:D_2_O (20:80) mixture at pH 7, resulting in isotope incorporation of ^1^H/^2^H at exchangeable sites at a ca. 20:80 ratio, and then lyophilized. Solutions used for crystallization were prepared by dissolving this lyophilized protein in buffers containing the same H_2_O:D_2_O ratio. All specified pH values below are corrected for the glass electrode isotope effect.^52^ For generating MPD-ub crystals, lyophilized ubiquitin was dissolved at 20 mg/ml concentration in buffer A (20 mM ammonium acetate, pH 4.3). Buffer B (50 mM citrate, pH 4.3) was mixed with 2-methyl-2,4-pentanediol (MPD) at a volume ratio of 40:60. 500 µL of this solution was deposited in the wells of sitting-drop crystallization plates. The protein solution (37µL) was mixed with the buffer B/MPD solution (10 µL) and deposited as sitting-drop in the crystallization plate (24-well plates with 1.5 mL reservoir volume). The plate was covered with sealing tape and kept at 4°C; crystals appeared after 1-2 weeks, and were filled into a 1.6 mm Agilent or 1.3 mm Bruker rotor using an ultracentrifuge device; caps were glued with epoxy glue to avoid dehydration. Cubic-PEG-ub crystals were prepared with the same equipment, but the protein was dissolved in buffer C (20 mM ammonium acetate pH 4.3) and mixed with buffer D (200 mM zinc acetate, 100 mM MES, pH 6.3, 20% w/v PEG 3350) and deposited as sitting-drop. The mother liquor reservoir was filled with 500 µL of buffer D. Crystals were obtained at 20 °C after circa 3-4 weeks.

## Solid-state NMR and data analysis

^15^N R_1ϱ_ relaxation measurements were performed on (i) a 600 MHz (14.1 T) Agilent VNMRS spectrometer (only BMCRD experiment) equipped with a 1.6 mm triple-resonance MAS probe tuned to ^1^H,^13^C,^15^N frequencies or (ii) a 600 MHz or (iii) a 950 MHz (22.3 T) Bruker Avance 2 spectrometer; the latter two were equipped with 1.3 mm MAS probes tuned to ^1^H,^13^C,^15^N frequencies with an auxiliary ^2^H coil. Resonance assignments of MPD-ub and cubic-PEG-ub have been reported elsewhere.^35^ BMCRD experiments at 600 MHz were performed at 39.5 kHz MAS, while BMCRD data at 950 MHz were collected at 50 kHz MAS. 600 MHz BMCRD data of MPD-ub have been reported before.^34^ NERRD data on both MPD-ub and cubic-PEG-ub were recorded on the 600 MHz (Bruker) spectrometer at a MAS frequency of 44.053 kHz. In order to enhance resolution, the BMCRD data on cubic-PEG-ub at 950 MHz were recorded as a series of proton-detected 3D hCONH-based experiments using a two-point approach with spin-lock periods of 1 and 50 ms). The 3D approach enabled us to resolve the otherwise overlapped I23 resonance. All other data were collected with 2D proton-detected hNH correlation experiments, shown in reference ^53^, using a series of relaxation delays (typically 8 points up to 125 ms). In all experiments, cross-polarization steps were used for transfer, with approximately 85 kHz ^1^H RF field, and a ^15^N RF field adjusted to the n=1 Hartmann-Hahn condition. Peak volumes in the individual 2D (3D) spectra were measured in NMRView (OneMoon Scientific), and relaxation rate constants were derived from monoexponential fits, using fitting routines written in python/numpy/scipy language. On-resonance *R*_1ρ_ relaxation rate constants were derived from the measured rate constants taking into account the resonance offset of the given cross-peak from the RF-field carrier Ω and the longitudinal relaxation-rate constant *R*_1_ as described elsewhere.^34^ This correction is fairly small, and visible primarily for low (<∼3 kHz) RF fields. Experiments at 950 MHz were measured twice, with two different carrier settings (112, 125 ppm), and the data reported for each residue correspond to the experiment with the carrier position closest to the resonance frequency. Error bars of relaxation rate constants were estimated from standard Monte Carlo simulations ^54^: the parameters obtained from fitting the experimental data set were used to generate a set of ca. 500 synthetic data sets, in which the peak intensities were randomly varied, relative to the back-calculated point, within three times the spectral noise level. These synthetic data sets were subsequently fitted in an identical way to the original data set, resulting in 500 fitted decay rate constants. The reported error bars correspond to the standard deviation of these rate constants.

Fits of a two-state exchange model as reported by Meiboom ^55^, to the BMCRD data were obtained from the program *relax*, version 4.0.^56^ For the fits shown in Figure 1 we chose five residues for which data of both crystals were available: I23 (only 950 MHz, as no 600 MHz data were available for cubic-PEG-ub) and V26, K27, T55, D58 (600 and 950 MHz). We have investigated how the choice of residues, and the choice of static magnetic field strengths influences the outcome of these fits; we find that using different subgroups of residues in the β-turn region, or including other residues in the fits where available (e.g. T22 in MPD-ub) does not significantly alter the results (see Supplementary Figure 5).

For the analysis of NERRD data we used numerical simulations, rather than the analytical equations (eq. 1). We prefer numerical simulations, because Eq. 1 reports accurately on the initial slope of relaxation decays, but fails to describe the multi-exponential behavior of the relaxation due to crystallite averaging. Experimentally determining initial slopes has been done, ^57^ but we instead choose (i) to use monoexponential fits of the experimental data and (ii) to interpret the resulting rate constants by using numerical simulations of decays, imitating the experimental setup as closely as possible. Details about the numerical simulations of the spin evolution and numerical fits are described in the Supporting Information.

## MD simulations

The 2-μs-long MD simulations of MPD-ub crystal (24 ubiquitin molecules) and cubic-PEG-ub crystal (48 ubiquitin molecules) recorded under Amber ff99SB*-ILDN force field ^58-60^ in the presence of SPC/E water^61^ are the same as reported by us previously,^35^ and described in detail in the Supplementary Methods. Analogous 2-μs-long trajectories have been recorded for MPD-ub crystals containing single-residue mutations E24A, G53A, K63A, E64A, or a negatively charged form of E24. For cubic-PEG-ub form, we have recorded the duplicate 2-μs trajectory of the wild-type ubiquitin crystal and the additional trajectories for crystals with K11A, K11A (chain A only), and G53A mutations. All alanine substitutions did not create any steric conflicts in the initial crystal coordinates. The compound statistics from each crystal trajectory is equivalent to 2 × 24 = 48 or 2 × 48 = 96 μs of ubiquitin dynamics. (Strictly speaking, many short trajectories are not equivalent to one long one, but in our case the simulation length is sufficient to sample the events of interest, i.e. conformational exchange and rocking). The trajectories exhibit a certain amount of drift, as has been discussed by us earlier;^35^ the drift is essentially a reflection of rocking motion that becomes progressively worse during the course of the simulations. In addition, we have also recorded a 1-μs-long trajectory of cubic-PEG-ub crystal, where C^α^ atoms within the secondary structure of ubiquitin have been restrained to their respective positions in the crystallographic structure by means of harmonic restraints with force constant 5 kcal/(mol·Å ^2^). The rocking motion was largely suppressed in this trajectory, leading to decreased formation of K11 _chainA_-D52_chainB_ salt bridge and lower content o f βII species. However, this trajectory featured a significant proportion of distinctive βII’ species, which appears to be an artefact. Therefore the restrained simulation was discontinued. As a control we have also recorded 2-μs-long trajectories of MPD-ub crystal and cubic-PEG-ub crystal (wild-type, K11A, K11A chain A only, and G53A) using newer ff14SB force field. ^62^ Finally, the previously reported solution trajectory of ubiquitin has been extended to 10 μs.

In processing the MD trajectories we used geometric criteria to identify hydrogen bonds (nitrogen-oxygen distance less than 3.2 Å, angle from 130° to 180°) and salt bridges (at least one nitrogen-oxygen distance less than 4Å, centroid distance 5Å). ^63,64^ Chemical shifts have been calculated by processing the MD coordinates using the program SHIFTX.^50^ This program has essentially the same accuracy as SHIFTX+ module from SHIFTX2,^65^ while relying on a simpler parameterization which nicely captures the dependence of ^15^N chemical shift on several essential structural variables. Given the limited accuracy of chemical shift prediction software, only the more pronounced modulation effects with Δδ(^15^N) on the order of 1 ppm can be reliably identified. Special tests have been performed to evaluate intermolecular contributions into chemical shifts; we have found that these contributions can be safely neglected for the problem at hand.

### Data availability

Solid-state NMR relaxation data and MD simulation frames can be obtained from the authors upon request.

## Acknowledgements

We thank Troels Schwarz-Linnet (University of Copenhagen) for helpful advice about the use of the program relax for data analysis, as well as Dr. Peixiang Ma, Jens D. Haller and Oleg Mikhailovskii and Manfred Burghammer for insightful discussions. This work was financially supported by the European Research Council (ERC-Stg-2012-311318-ProtDyn2Function), the French Research Agency ANR (ANR 10-PDOC-011-01), as well as Commissariat à l’énérgie atomique et aux énérgies alternatives (CEA), Centre National de la Recherche Scientifique (CNRS) and Université Grenoble Alpes. This work used the platforms of the Grenoble Instruct Center (ISBG; UMS 3518 CNRS-CEA-UJF-EMBL) with support from FRISBI (ANR-10-INSB-05-02) and GRAL (ANR-10-LABX-49-01) within the Grenoble Partnership for Structural Biology (PSB). S. A. I., O. N. R. and N. R. S. acknowledge Russian Science Foundation for financial support of the MD simulation component of this study (RSF grant 15-14-20038). N. C. is supported by a fellowship from the Fondation France Alzheimer.

## Author contributions

V. K. and P. S. performed and analyzed NMR experiments. A. H. and I. A. prepared protein samples and crystals. P. S., A. H., J. W., N. C., J.-P. C. and A. S. crystallized ubiquitin in different crystal forms or characterized crystal samples. S. A. I., O. N. R., Y. X., T. Y. and N. R. S. performed and analyzed MD simulations. P. S. and N. R. S. designed the research. P. S. and N. R. S. wrote the manuscript with contributions from other authors.

